# Viral infection collapses intracytoplasmic membrane integrity and autotrophic metabolism in ammonia-oxidizing *Nitrosomonas europaea*

**DOI:** 10.64898/2025.12.21.695702

**Authors:** J. Papendorf, D. K. Ngugi, P. Büsing, N. Reimann, J. Wittmann, S. Peter, R. L. Hahnke, S. Kirstein, B. Bunk, M. Rohde, M. Müsken, M. Neumann-Schaal, M. Pester

## Abstract

Ammonia-oxidizing bacteria (AOB) catalyze the first and rate-limiting step of nitrification. They are essential for nitrogen cycling in engineered and natural environments, yet little is known about their viruses or the consequences of phage infection for host physiology. Here, we report the isolation and characterization of a novel lytic bacteriophage, vB_NeuP-Nir1 (Nir1), infecting the model AOB *Nitrosomonas europaea*. Phage Nir1 was highly virulent, ceased ammonia oxidation within hours, and caused complete lysis of host populations even at multiplicities of infection as low as 10^−6^. Electron microscopy revealed drastic host cell remodeling during infection, including pronounced cell bloating and large-scale disintegration of intracytoplasmic membranes. Integrated transcriptomic and metabolomic analyses showed that loss of these ATP and reducing equivalent generating membrane systems was accompanied by signatures of compromised lipid homeostasis and collapse of autotrophic CO₂ fixation. In parallel, Nir1 infection induced metabolic rewiring of the host, including upregulation of uptake systems for nucleic acids, amino acids, and small organic compounds, increased expression of iron acquisition and putative iron-dependent respiratory components, as well as accumulation of metabolites associated with membrane breakdown and stabilization of viral DNA. Together, these results provide the first detailed mechanistic insight into phage-induced host modulation in a chemolithoautotrophic nitrifier. Our study establishes the Nir1–*N. europaea* system as a model for investigating virus–host interactions in AOB and lays the foundation for assessing the role of phages in shaping nitrification and nitrogen cycling in engineered and natural ecosystems.

## Introduction

The nearly ubiquitous process of biological nitrification is an important part of the global nitrogen cycle. It describes the aerobic conversion of ammonia via nitrite to nitrate and extends from marine over terrestrial to man-made ecosystems [1]. In ammonia-rich environments such as agricultural soils, the transformation of ammonia to leachable nitrate facilitates fertilizer loss and can ultimately lead to eutrophication of neighboring aquatic ecosystems [2]. Additionally, nitrification releases the reactive nitrogen species NO, HONO, NO_2_ and N_2_O to the atmosphere [3–5]. Contrasting these undesirable effects, nitrification is an essential part of biological nitrogen removal from wastewater [6], which enables discharge of the effluent without causing eutrophication of affected water bodies and harm to aquatic life.

Depending on the physicochemical parameters of given habitat and niche adaptation, different ammonia-oxidizing microorganisms (AOM) are dominant and perform the first and rate-limiting step of nitrification [7–10]. Ammonia-oxidizing bacteria (AOB) are typically abundant in ammonia-rich environments [11, 12]. In contrast, different lineages of ammonia-oxidizing archaea (AOA) vary in their substrate concentration preference across a broad range of ammonia concentrations, including a strong predominance in oligotrophic, low-ammonia environments [9]. AOB and AOA cooperate with nitrite-oxidizing bacteria (NOB) for further conversion of AOM-derived nitrite to nitrate [13]. Owing to their ability of complete ammonia oxidation to nitrate, a recently discovered third group of AOM was named Comammox [14, 15]. Similar to part of AOA, Comammox have been associated with an oligotrophic lifestyle [16] and found in low-ammonia environments [17, 18].

Representatives of all three AOM groups have been detected in wastewater treatment plants (WWTP) [19–22]. Nevertheless, AOB often dominate the WWTP AOM-community at elevated ammonia concentrations with most representatives being affiliated with the genus *Nitrosomonas* [20, 23, 24]. *Nitrosomonas europaea* is common in WWTPs and represents a model organism for AOB [11]. Much of what is known about the physiology, biochemistry, and molecular biology of AOB is based on studies of this chemolithoautotrophic Betaproteobacterium [25]. It tolerates high concentrations of ammonia (up to 100 mM) and nitrite (up to 25 mM) [25]. Due to the toxicity of intermediates and products (i.e. hydroxylamine, NO and nitrite) of its energy metabolism, the respective enzymatic machinery is located mainly in the membrane coupled to a release of metabolites into the periplasmic space [26]. This explains, why AOB including *N. europaea* are characterized by a high degree of intracytoplasmic membrane stacks to increase the surface for their energy metabolism [27]. *N. europaea* is typically growing autotrophically by utilizing the Calvin cycle, although reports exist where *N. europaea* grew as a facultative chemolithoheteroph with fructose or pyruvate as resource for its anabolic pathways [28]. The sum of this knowledge makes *N. europaea* a highly suitable model organism to study principles of virus-host interactions for AOM in high-ammonia settings.

Viruses that infect prokaryotes have first been described more than 100 years ago. Today, we know that they are omnipresent and abundant in the environment [29–31] and man-made systems like WWTPs [32]. It has been postulated and partly already shown that they play an important role in shaping microbial communities and associated carbon cycling [33–36]. However, little is known about the impact of viruses on nitrogen cycling and especially AOM. Regardless of clade, growth of AOM including *N. europaea* is slow and generally only occurs in liquid cultures or takes multiple weeks on solidified medium. This renders typical microbiological and virus-associated methods futile and demands for alternative techniques. It also explains why virus-host interactions have been extremely little studied for these chemolithoautotrophic microorganisms and mainly relate to descriptions in metagenomic datasets [37–39]. So far, only two AOM-infecting viruses, a chronically AOA-infecting virus [40] and a lytic AOB-infecting bacteriophage [41] have been isolated. However, studies targeting viral host modulation of AOM are completely lacking.

In this study, we isolated a bacteriophage infecting *N. europaea* from a wastewater treatment. Using infection assays, electron microscopy, transcriptomics, and metabolomics, we provide the first integrated analysis of phage-induced host modulation in a chemolithoautotrophic nitrifier. Phage infection rapidly halted ammonia oxidation, triggered disintegration of intracytoplasmic membranes, disrupted lipid homeostasis and CO_2_ fixation, and induced extensive metabolic rewiring to support viral replication. Our results extend knowledge on viral host modulation beyond well-studied heterotrophic [42] or phototrophic [43] model prokaryotes to chemolithoautotrophs catalyzing a key step in the N cycle.

## Material and Methods

### Isolation and genome analysis of phage vB_NeuP-Nir1

*N. europaea* strain Nm50^T^ was grown for 19 days in 100 mL DSMZ medium 1583 (Supplementary Material and Methods) at 28°C in the dark without agitation (https://mediadive.dsmz.de, Koblitz et al., 2023). Ninety-nine milliliters of the bacterial culture were then concentrated via centrifugation for 15 min at 14,334×*g* and 15°C. The bacterial pellet was resuspended in 1 mL of the supernatant. Initial infection of the bacterial culture was performed by the addition of 8 mL fresh DSMZ medium 1583 and 1 mL 0.22-µm-filtered (Corning®, PES, Corning Inc., Corning, USA) activated sludge obtained on December 2^nd^, 2019 from the WWTP Steinhof located in Braunschweig, Lower Saxony, Germany (52°19’05.9"N, 10°26’45.1"E).

Positive infection was indicated by the absence of ammonia oxidation activity, which results in pH drop of the medium and can be monitored by the included pH indicator cresol red. Isolation of the bacteriophage was performed by repeated serial dilution of 1 mL of infected host cultures in 10 mL culture tubes containing 8 mL fresh DSMZ medium 1583 and 1 mL of the host strain grown for ca. 2 weeks in DSMZ medium 1583 before infection. Purity of the isolated phage was confirmed by electron microscopy and sequencing of the phage genome. For the latter, short-read Illumina and long-read ONT sequencing were combined. For Illumina sequencing, bacterial debris was removed by centrifugation for 15 min at 14,334×*g* and 15°C. Concentration of phage lysate (3.5 L) was achieved by repeated centrifugation in 33 mL aliquots for 2 h at 36,358×*g* and 4°C. The majority of supernatant was removed by pipetting and pellets resuspended in residual supernatant at 120 rpm on an orbital shaker at RT for 1 h. Concentrated viral lysates were then pooled, resulting in a final volume of 9 mL. For ONT sequencing, bacterial debris was removed by 0.2-µm-filtration (Filtropur S 0.2, PES, Sarstedt, Nürmbrecht, Germany). Concentration of phage lysate was achieved by centrifuging 2×20 mL phage lysate in Macrosep 30 kDa centrifugal devices (Pall, Crailsheim, Germany) for 30 min at 1,000×*g*. For both, Illumina and ONT sequencing, isolation of phage DNA from concentrated lysates was performed using the Phage DNA Isolation Kit (Norgen Biotek Corp, Thorold, Canada) according to the manufacturer’s protocol including optional steps of Proteinase K and DNase treatment.

ONT sequencing was performed on a MinION device using the ligation sequencing kit SQK-LSK114 according to the protocol but omitting the addition of DNA Control Sample (DCS) and the Flongle Flow Cell R10.4.1 (Oxford Nanopore Technologies, Oxford, United Kingdom). Libraries for Illumina sequencing were prepared as previously described [44]. Illumina sequencing was performed on the MiSeq System (Illumina, San Diego, USA) with 2×300 bp read length. Assembly of SUP (super accurate) base-called reads from ONT sequencing was done using Flye [45, 46] version 2.9. In Geneious Prime version 2025.0.2, reads produced via Illumina sequencing were Q30-trimmed and filtered for a minimal length of 10 nucleotides using BBDuk (part of BBMap suite) v38.84 [47]. Reads were then mapped to the assembled contig using Bowtie 2 version 2.4.5 [48] to correct ambiguities from ONT sequencing. Gene annotation was done with Pharokka version 1.7.5 [49]. Further annotation of hypothetical genes was done with Phold version 0.2.0 [50]. One CDS had to be manually corrected to begin with a start codon. A circular plot of the viral genome was created using the plot option in Phold [50]. Phylogenomic reconstruction was done in ViPTree (Viral Proteomic Tree) server version 4.0 [51]. Tree calculation was done using default settings but with additional gene functional prediction using GHOSTX. Calculation of intergenomic similarities between phages was performed using the web servers of taxMyPhage v3.3.6 [52] and of VIRIDIC [53] with default parameter settings.

### Electron microscopy

Fixation of bacteria was done in a stepwise manner in a concentrated bacterial suspension in the growth medium. First, 25% glutaraldehyde (final concentration 2%) was added and incubated for 30 min, followed by addition of 25% paraformaldehyde (final concentration 5%). SEM and TEM sample preparation for ultrathin sections was done as previously described [54, 55] but making use of an automated critical point dryer CPD300 (Leica Microsystems, Wetzlar, Germany). For SEM preparation, in addition to acetone dehydration, ethanol dehydration of the samples was performed without prior washing steps. Fixation of the sample on the cover slip was performed overnight due to a low bacterial concentration. Visualisation of phages and flagella was done according to [56] using 0.5% uranyl acetate for negative staining of bacteria and 2% for phages and flagella. The latter were separated by strong vortexing to disconnect flagella from bacteria and in a two-step centrifugation procedure. First to separate bacteria and second to concentrate flagella (16,060×*g*, 30 min at RT).

### Host range experiments

Bacterial strains were grown in DSMZ medium 1583 at 28°C in the dark without agitation. Actively growing cultures (n=3) of AOB *Nitrosomonas eutropha* C-91^T^ (=DSM 101675), *Nitrosomonas communis* Nm2^T^ (=DSM 28436), *Nitrosomonas nitrosa* Nm90^T^ (=DSM 28438) and *Nitrosospira multiformis* C-71^T^ (=DSM 101674) were amended with 0.2-µm-filtered viral lysate of Nir1. Strain Nm50^T^ was used as positive control to confirm infectivity of the virions. After three and seven days, all cultures were investigated regarding color change of the pH indicator as an indicator for ammonia-oxidizing activity and in addition by phase contrast microscopy regarding bloated and lysed cells.

### Defined infection experiments with different MOIs

Infection experiments were run in four replicates for multiplicities of infection (MOIs) 9.2×10^−1^, 1.6×10^−1^, 8.6×10^−2^ and 1.4×10^−2^, and in duplicates for MOIs 1.3×10^−4^ and 1.2×10^−6^. Virion concentrations were determined by counting SYBR-Gold (Invitrogen, Life Technologies Corporation, Eugene, USA) stained virions on a 0.02 µm pore-size Anodisc aluminium oxide filter membrane (Cytiva, Marlborough, MA, USA) using 100 randomly selected fields of 20×20 µm or 400 fields of 10×10 µm. Phage staining was performed as described by [40] with minor modifications (Supplementary Material and Methods).

We followed bacterial growth and nitrite production starting 43 h before and ending 121 h after addition of virions. For the experiment done in duplicates, infection was monitored only until 29 h post infection. Host populations were followed by flow cytometry. Sample preparation of bacterial populations consisted of filtration through a 10 µm cell-sieve (CellTrics™, Sysmex Partec GmbH, Görlitz, Germany) to remove precipitates of the medium followed by bacterial staining for 10 min in the dark using 1 µL 500 µM SYTO-9 (Invitrogen) added to 999 µL bacterial culture. Bacterial concentrations were determined using a CytoFlex S benchtop flow cytometer (Beckman Coulter, Brea, USA) and co-analysis with 4 µm CountBright Plus Absolute Counting Beads (Invitrogen) as internal standard. A total of 50,000 or 100,000 events were observed per sample using a threshold of 4000 in the SSC-H channel and adjusted gain settings (forward scatter FSC=120, side scatter SSC=700, SYTO-9=1750). Events representing bacterial cells and counting beads were captured in gates created in SYTO-9 vs. SSC and FSC vs. SYTO-9 plots, respectively. Nitrite concentrations were quantified on a S150 Ion Chromatography System (Sykam Chromatographie Vertriebs GmbH), equipped with a SykroGel AX300 column, using 0.025 mM NaSCN and 4 mM Na_2_CO_3_ as eluent and a flow rate of 1 mL min^−1^.

### Transcriptome response of Nm50^T^ to Nir1-infection

Isolation of genomic DNA and reconstruction of a closed reference genome of the host Nm50^T^ was performed as described in Supplementary Material and Methods. Bacterial and viral concentrations were quantified as described above. A culture of strain Nm50^T^ was grown statically in a 5 L Erlenmeyer flask for 11 days at 28°C in the dark. The bacterial culture was split into six 1 L Erlenmeyer flasks, each containing 290 mL of the culture. Three of the replicates were amended with 7.38 mL phage lysate of Nir1 (MOI=4.1) and gently mixed, the remaining three replicates served as control. Infected and non-infected cultures sampled directly after addition of virions and after 120 min of static incubation at 28°C in the dark. For each timepoint and replicate, 140 mL of culture were collected on ice and directly centrifuged in 250 mL PPCO centrifuge bottles (Nalge Nunc International Corporation, Rochester, USA) for 30 min at 4°C and 9,000×*g*. The supernatant was carefully decanted and the pellet was directly resuspended in 600 µL lysis buffer of the AllPrep PowerViral DNA/RNA Kit (Qiagen, Hilden, Germany). RNA extraction was performed immediately after sampling as described by the manufacturer. Extracted nucleic acids were eluted in molecular grade, nuclease-free water (Thermo Fisher) and directly frozen at −80°C until further analysis. Presence of RNA and DNA was confirmed via 1% agarose gel electrophoresis (130 V, 30 min) using 1% Wide Range agarose (SERVA Electrophoresis GmbH, Heidelberg, Germany) in 1× TAE buffer (Carl Roth GmbH + Co. KG, Karlsruhe, Germany). DNA was then removed using DNase I treatment (rDNase I, DNA-*free*™, Invitrogen). In brief, 9.6 µL of 10× DNase buffer was added to 100 µL of each sample followed by addition of 1.5 µL of rDNase (2 U µL^−1^). Samples were incubated for 30 min at 37°C. After initial incubation, one additional microliter of DNase was added and samples incubated again for 30 min at 37°C. Inactivation of DNase was achieved by addition of 19.2 µL DNase inactivation reagent followed by thorough mixing and incubation for 5 min at RT. Samples were then centrifuged for 1.5 min at 10,000×*g* at RT. Concentration of RNA was determined using the Qubit™ RNA HS Assay Kit (Invitrogen) according to the manufacturer’s protocol. Removal of genomic DNA was confirmed via 16S rRNA gene PCR (35 cycles) using general bacterial primers 27-f (5’-AGAGTTTGATYMTGGCTC-3’) and 1492-r (5’- GGYTACCTTGTTACGACTT-3’) at an annealing temperature of 52°C, followed by gel electrophoresis. Absence of specific PCR products was taken as a sign of efficient DNA removal.

Next, 50 µL of each sample were concentrated to 15 µL using the RNA Clean & Concentrator™-5 kit (Zymo Research, Irvine, USA) according to the manufacturer’s protocol. Concentration of RNA was determined using the Qubit™ RNA HS Assay Kit (Invitrogen) according to the manufacturer’s protocol. Fourteen microliters of each sample were then depleted for rRNA using the Ribo-off rRNA Depletion Kit for Bacteria (Vazyme biotech, Nanjing, PRC), according to the manufacturer’s protocol. Elution was performed in 8 µL molecular grade, nuclease-free water. Seven microliters of each sample were prepared for Illumina sequencing using the TruSeq Stranded mRNA kit (Illumina) according to the manufacturer’s protocol. Quality of libraries was confirmed via nucleic acid fragment analysis using a Femto Pulse System (Agilent Technologies, Inc., Santa Clara, USA). Sequencing was performed on a NextSeq 2000 Sequencing System (Illumina) using 2×150 bp. Reads were Q30-trimmed and filtered to a minimal length of 100 nucleotides using BBDuk (part of BBMap suite) v37.62 [47] followed by discarding singletons. Reads were then mapped to host CDS and phage CDS using BBMap v37.62 [47]. Differential expression of host genes was performed in R v4.4.1 (https://cran.r-project.org/) using the DESeq2 package [57] v1.46.0. Visualization of differentially expressed genes was performed using the EnhancedVolcano package v1.24.0 (https://github.com/kevinblighe/EnhancedVolcano) in R v4.4.1.

### Metabolome response of Nm50^T^ to Nir1-infection

Bacterial and viral concentrations were quantified as described above. Two bacterial cultures of Nm50^T^ were grown statically in 5 L Erlenmeyer flasks for 12 days at 28°C in the dark and mixed together directly prior to the experiment. From this homogenized host culture, 290 mL each were distributed to twelve 1 L Erlenmeyer flasks. Six replicates served as uninfected control and six replicates were amended with 11.4 mL phage lysate or Nir1 (MOI=3.44) for the infection experiment. Sampling took place directly at the start of each treatment and after 120 min of static incubation at 28°C in the dark. Since taking samples from six replicates per treatment takes a short time period including first phage activity, we defined the timepoint at the start of the infection experiment as early response and the timepoint after 120 min as late response. For each timepoint and replicate, 140 mL of culture were centrifuged in 250 mL PPCO centrifuge bottles (Nalge Nunc International Corporation) for 30 min at 4°C and 9,000×*g*. The supernatant was carefully decanted, the pellet resuspended in 1 mL fresh DSMZ medium 1583 and centrifuged again for 15 min at 4°C and 9,000×*g*. The supernatant was removed with a pipette, the biomass immediately frozen in liquid nitrogen and stored at −80°C.

For GC-MS analysis, biomass was extracted by adding 250 µL methanol, spiked with 0.5% ribitol (0.2 mg mL^−1^), for 15 min at 70°C in an ultrasonic bath. Samples were incubated for 2 min on ice before 250 µL water were added. Samples were mixed vigorously. Subsequently, 300 µL dichloromethane were added and samples were mixed again. Following a centrifugation step (5 min, 10,000×*g*), the polar phase was collected and 400 µL were dried under vacuum. Samples were derivatized and measured as described earlier on an Agilent GC-MSD system (7890B coupled to a 5977 GC) equipped with a high-efficiency source (HES) and a Gerstel RTC system [58]. In brief, a two-step derivatization with a methoxyamine hydrochloride solution (20 mg mL^−1^ in pyridine) and *N*-methyl-*N*-(trimethylsilyl)-trifluoracetamide was automatically performed followed by separation on an Agilent VF-5ms column and analysis in scan mode. Data analysis of intracellular metabolites was performed as previously described [59, 60].

### Diagnostic PCR screening for Nir1

In the WWTP Steinhof, secondary clarifiers are fed by three effluents from corresponding aeration basins. Each effluent was sampled once per month for twelve consecutive months (February 2024 – January 2025). Samples were taken with a custom rod sampler which was flushed three times in the wastewater stream before taking samples. Using serological pipettes, 25 mL of wastewater were transferred to 50 mL conical polypropylene centrifuge tubes (TPP Techno Plastic Products AG, Trasadingen, Switzerland), transported to the laboratory and frozen at −20°C until analysis. Samples were vortexed for 30 sec after thawing since the flocculated activated sludge had settled. DNA extraction was performed on a 200 µL subset of activated sludge from each sample using the AllPrep PowerViral DNA/RNA Kit (Qiagen) according to the manufacturer’s protocol, using beat beating to ensure complete lysis.

Primer-BLAST [61] was used to design the PCR primer pair cap3f (5’-TGCTACGGAGGCTAAAGCTCG-3’) and cap3r (5’-CTACTGCATCACCACGCAGC-3’) targeting a 170 bp long region of the gene encoding the major head protein of Nir1. A specificity check via Primer-BLAST against the NCBI nucleotide collection (nt) database detected only the highly similar genome of phage ΦNF-1 (OL634959.1) as additional potential target with a single nucleotide mismatch for one of the two primers. PCR was performed with 60 ng template, 0.4 µL of each primer (10 µM), 10 µL 2× DreamTaq Green PCR Master Mix (Thermo Fisher Scientific Baltics UAB, Vilnius, Lithuania) and a total volume of 20 µL on a Biometra TOne (Analytik Jena GmbH+Co. KG, Jena, Germany) thermocycler. Genomic DNA (0.121 ng) of Nir1 served as positive control. Molecular grade, nuclease-free water served as negative control. PCR was performed with 95°C initial denaturation, 35 cycles of 95°C, 60°C, 72°C each for 30 sec, and 5 min of final elongation at 72°C.

### Statistics

Graphs and Kruskal Wallis nonparametric tests with Dunn’s multiple comparisons tests were performed using the Graph function of BioRender.

## Results

### *N. europaea*-infecting phage Nir1 persists year-round in a German WWTP

We isolated the lytic bacteriophage vB_NeuP-Nir1 (=Nir1) infecting the AOB *Nitrosomonas europaea* DSM 28437 (=Nm50^T^) by amending an actively growing host culture with 0.2-µm-filtered wastewater followed by serial dilution. Electron microscopy revealed a T7-like podoviral shape of virions with a capsid size of 60-65 nm in diameter and a short (10-15 nm) tail (Fig. 1a). Combining long (ONT) and short (Illumina) read sequencing, we assembled its 41,212 bp-long genome (642-fold mean coverage, 99.1% Q30) and were able to annotate 29 of 61 predicted CDS (Fig. 2a). A ViP-Tree-based [51] comparison of genome-wide similarities computed by tBLASTx revealed closest relatedness to bacteriophages of the former family *Autographiviridae*, recently promoted to the novel order *Autographivirales* [62], infecting bacteria of the phylum *Pseudomonadota* (Fig. 2b). According to the Virus Intergenomic Distance Calculator VIRIDIC [53], Nir1 possesses 90.56% intergenomic similarity to ΦNF-1, the first described phage infecting multiple *Nitrosomonas* species [41], but only 4.62% similarity to the next closest known related phage vB_SnaP-R1 (=SnaR1), infecting heterotrophic *Sphaerotilus natans* strain DSM 6575 [63]. With 90.8% calculated similarity of Nir1 to ΦNF-1, results from the taxonomic assignment tool taxMyPhage [52] were highly similar. Following the demarcation thresholds of the ICTV [64], Nir1 was determined by taxMyPhage and VIRIDIC as a novel species in the recently established genus *Catalonvirus*, currently containing solely species ΦNF-1. Nir1 was deposited at DSMZ under the accession number DSM 111086.

**Figure 1.**
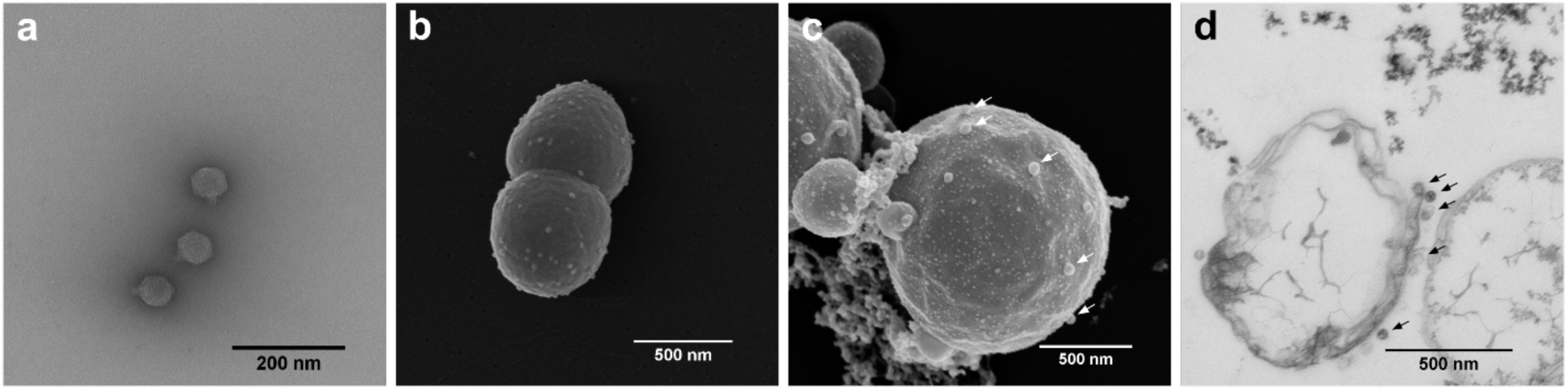
Morphology of bacteriophage vB_NeuP-Nir1 (Nir1) as well as uninfected and Nir1-infected cells of *Nitrosomonas europaea* Nm50^T^. a) TEM of podoviral bacteriophage Nir1, b) SEM of uninfected *N. europaea* Nm50^T^ cells showing typical morphology, c) SEM of Nir1-infected *N. europaea* Nm50^T^ cells showing a bloated morphology and vesicle production, white arrows mark representative virions d) TEM of thin-sliced Nir1-infected *N. europaea* Nm50^T^ cells showing disintegrated intracellular membranes, black arrows mark representative virions.

**Figure 2.**
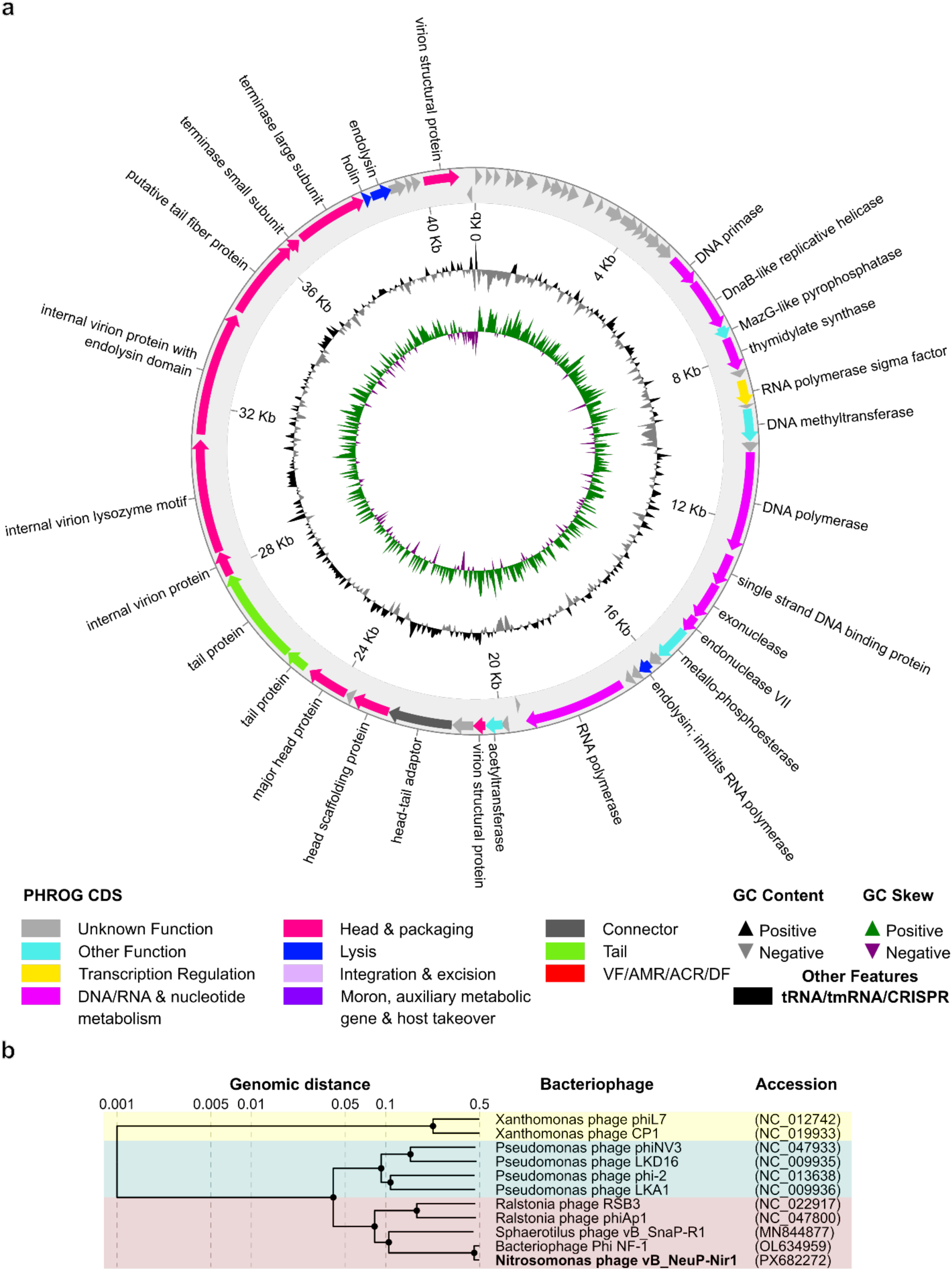
Genome characteristics and phylogeny of bacteriophage vB_NeuP-Nir1. a) Genome organization and annotated genes of bacteriophage vB_NeuP-Nir1, b) BIONJ-based [102] user-generated viral proteomic tree created from genomic distances derived from normalized tBLASTx scores [51]. Genomic nucleotide sequences of vB_SnaP-R1, ΦNF-1 and vB_NeuP-Nir1 were uploaded to the ViPTree server using dsDNA as nucleic acid type and prokaryote as host category of reference viruses. Close relatives and an outgroup were manually selected from the complete sequence list provided with the calculated tree and used to create a user-generated tree.

While Nir1 was isolated in northern Germany, ΦNF-1 was isolated from a WWTP located in Catalonia, more than 1,200 km apart. We further followed the presence of Nir1 in its original habitat, the WWTP Steinhof, throughout the year using a diagnostic PCR specifically designed to target the gene encoding its major head protein. Nir1-specific PCR products were detectable year-round but were most pronounced from July to October (Supplementary Fig. 1).

### Nir1 is highly virulent and severely alters host morphology

Upon infection with Nir1, cells of Nm50^T^ undergo a drastic morphological change from short coccoid rods (Fig. 1b) towards a swollen spherical phenotype, sporadically with vesicular membrane extrusions (Fig. 1c). Occasionally, aggregation of cells occurred in infected cultures. To investigate intracellular changes of infected bloated host cells, we performed transmission electron microscopy of sliced cells. We observed that intracytoplasmic membranes as characteristic for bacteria of the genus *Nitrosomonas* [65] and other nitrifiers [27] were disintegrated. Virions were not detected within infected bloated host cells (Fig. 1d) unless infection deemed incomplete, indicating that progeny is generally efficiently released. To test the ability of Nir1 to infect alternative hosts, we screened four described AOB species of the genera *Nitrosomonas* and *Nitrosospira*, including all known host species of ΦNF-1 [41]. Only a minor fraction of *N. nitrosa* DSM 28438 (=Nm90^T^) cells showed typical signs of infections (i.e., hollow or bloated cells) while all other screened hosts were infection negative. To analyze the virulence of Nir1, infection experiments with defined virus-to-host ratios were performed in liquid cultures with MOIs ranging from 9.2×10^−1^ down to 1.2×10^−6^. MOI is defined here as the ratio of phage particles to host cells at the moment of inoculation. For the highest MOI tested, onset of lysis was detectable ca. 3 h post infection, which coincided with a stop of nitrite production as a proxy for metabolic activity. A decrease of MOI gradually slowed the infection process and delayed the stop of metabolic activity. However, also under the lowest MOI tested (1.2×10^−6^) metabolic activity ceased after 25.5 h and complete lysis of the host population was observed 29 h post infection, showing the high virulence of Nir1 (Fig. 3).

**Figure 3.**
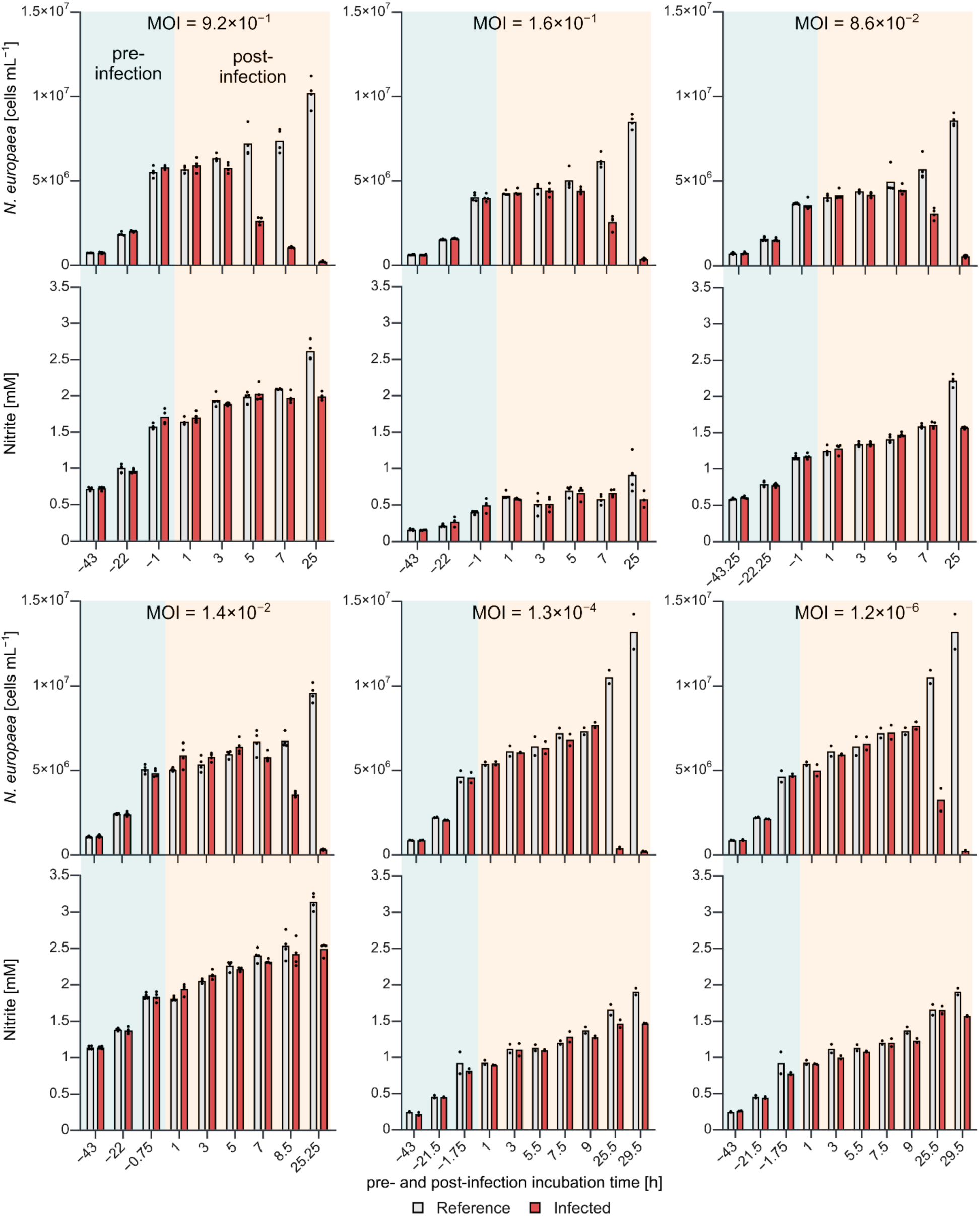
Infection experiments of *N. europaea* Nm50^T^ with phage vB_NeuP-Nir1 under different multiplicities of infection (MOI). Host cell numbers and nitrite concentration are shown as proxies for bacterial growth and activity, respectively (n=4; except for MOIs 1.3×10^-4^ and 1.2×10^-6^, n=2).

### Nir1 infection triggers overexpression of genes encoding membrane-associated proteins

We next analyzed host gene expression during the early and late infection process. This was paralleled by de-novo sequencing of the genome of Nm50^T^ to provide a high-quality and closed reference genome. Genes were considered significantly (adjusted *p*-value <0.05) differentially expressed if log_2_ fold changes (LFC) were above 1 or below −1. Comparison of early infected against non-infected cells did not reveal significant differential expression of genes. Cells from late sampled controls showed a significant differential expression of nine genes compared to early non-infected cells. All of these genes were downregulated with LFCs between 1.01 and 1.71 and coded for a phosphate transporter, a putative thiosulfate transporter, four RuBisCO (ribulose-1,6-bisphosphate carboxylase/oxygenase) associated gene products and three hypothetical proteins.

Nir1-infected cells during the late infection process showed significant transcriptional upregulation of 142 genes compared to late non-infected cells but no downregulation (Fig. 4, Supplementary Table 1). Two genes coding for multicopper oxidases showed high overexpression (LFC of 6.16 and 6.14, respectively), with four additional genes coding for redox-active proteins, including cytochrome c, being overexpressed as well (LFC 1.19-1.61). Further transcriptionally highly upregulated genes (LFCs 5.05-6.27) encoded an amino acid permease, a urea carboxylase and three urea amidolyase gene products. With LFCs ranging between 1.35 and 3.93, we detected overexpression of nine genes involved in iron uptake or associated with low iron availability.

**Figure 4.**
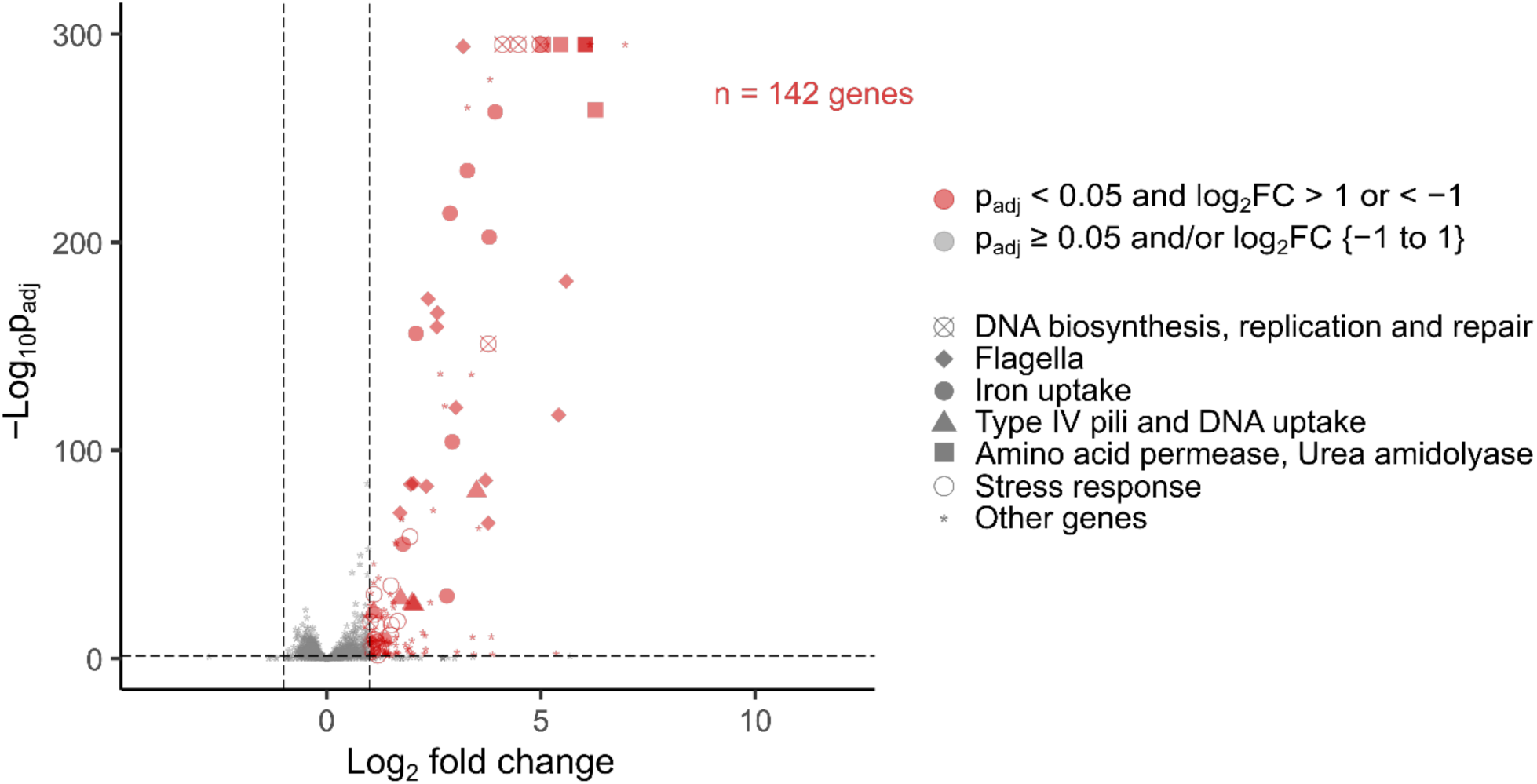
Differentially expressed genes of Nir1-infected *N. europaea* Nm50^T^ in comparison to uninfected controls. Samples were taken at a timepoint corresponding to the late infection process (120 min post infection, n=3). Log_2_-fold change (LFC) cut-offs were set to 1 and −1 for up- and downregulated genes, respectively, and are represented by dashed grey vertical lines. Adjusted *p*-values (*p*_adj_) are shown as −Log_10_p_adj_. The significance cut-off for adjusted *p*-values of 0.05 is represented by a dashed grey horizontal line. Significantly differentially expressed genes are marked in red, all other genes are shown in grey.

Notably, two genes coding for flagellar transcriptional activators (*flhD* and *flhC*) with an adjacent gene coding for a ComEC/Rec2 protein (LFC 3.50) and a gene cluster consisting of at least 12 flagellar genes (*flgA*, *flgB*, *flgC*, *flgD*, *flgE*, *flgF*, *flgG*, *flgH*, *flgI*, *flgJ*, *flgK* and *flgL*) were highly upregulated as well (LFCs 1.01-5.59). Furthermore, we detected four overexpressed genes (LFCs 1.10-2.04) involved in Type IV pilus biogenesis (*pilP*, *pilO*, *pilN* and p*ilQ*). We also observed the overexpression of one gene coding for carbon starvation protein A (LFC 2.65) and, directly adjacent, one gene coding for an YbdD/YjiX family protein (LFC 2.80). Other categories of overexpressed genes include stress response, DNA synthesis, replication and repair, transcriptional regulators, ribosomal proteins and cell wall remodeling (Supplementary Table 1). In addition, 27 upregulated genes coded for proteins of unknown function and eleven for transposases. Investigation of infected cells via electron microscopy did not reveal enhanced production of cellular appendices such as flagella or pili but confirmed onset of bloating and vesicle formation 3 hours post infection.

### Nir1 infection increases membrane-associated metabolite levels

To elucidate how infections of Nir1 impact the central metabolism of Nm50^T^, we analyzed changes in its respective metabolite profile during the early and late infection process in a similarly fashioned infection experiment as done for the transcriptomic responses. In total, profiles of 32 metabolites could be detected (Fig. 5) and were mapped via assigned K numbers to KEGG metabolic pathways. Concentrations of metabolites associated with lipid- and fatty acid metabolism were strongly and significantly elevated in Nir1-infected cultures compared to their controls. This effect was specifically pronounced for glycerophosphoglycerol and 3-hydroxybutanoate, respectively. We also detected strong and significant increases of the fatty acids hexadecanoate and hexadec-9-enoate, further supporting changes in membrane-associated metabolites. A second group of significantly altered metabolite profiles concerned the core central metabolism and encompassed pyruvate, 3-phosphoglycerate, malate, succinate, glyoxylate, and glycolate. While there was no common trend of up- or downregulation, the observed changes indicate altered routes of carbon flow within the Calvin cycle and the biosynthesis-supporting citric acid cycle and gluconeogenesis of these autotrophic bacteria. The third major group of altered metabolite profiles concerned various amino acids, as would be expected from a shifted amino acid demand of proteins involved in the infection process including phage-specific proteins. In addition, the polyamine putrescine had significantly higher cellular concentrations in infected cells at the late infection process timepoint. Further metabolites with altered concentrations profiles concerned the sugars glucose, fructose, and galactose as well as AMP as a central metabolite directly involved in energy metabolism.

**Figure 5.**
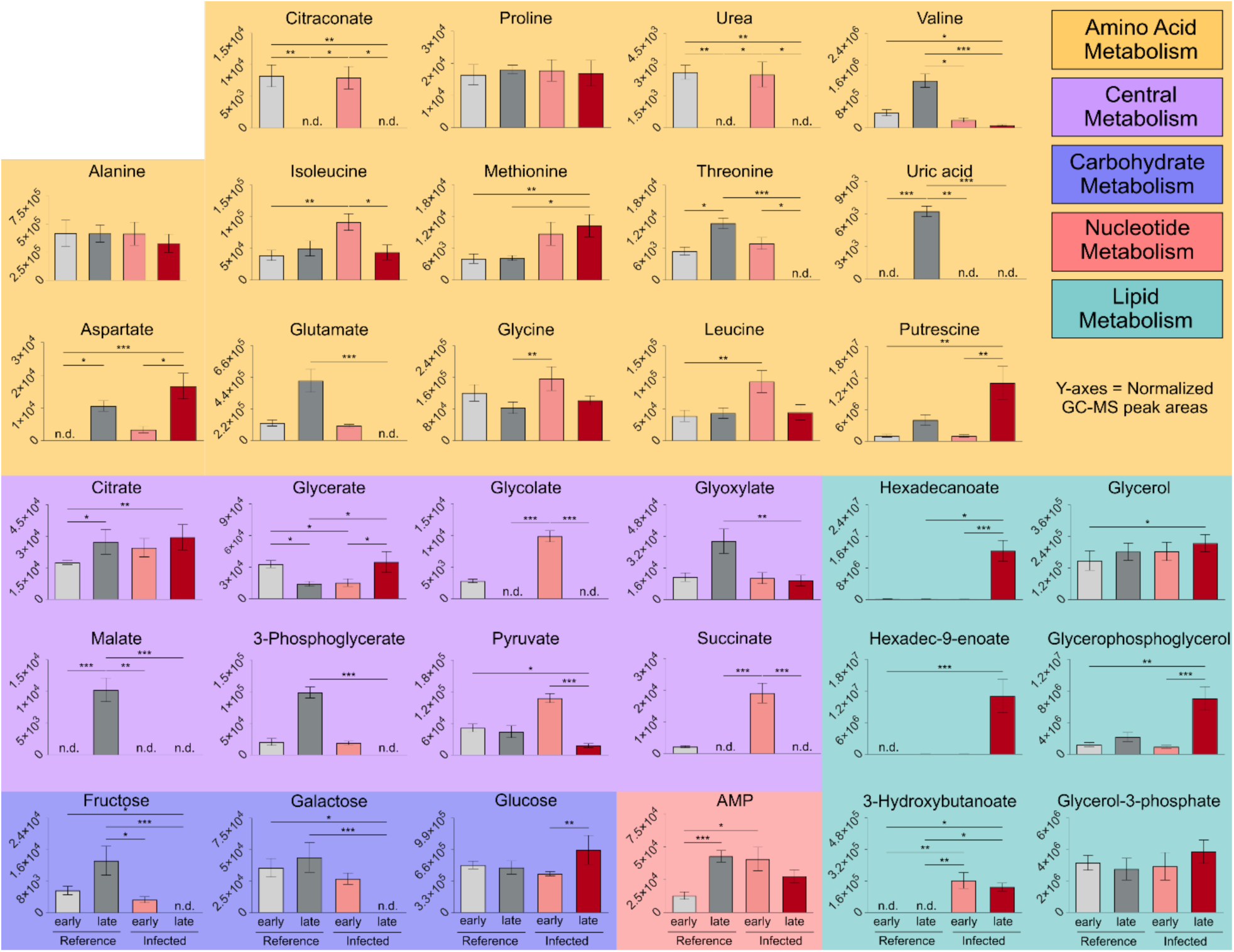
Metabolite profiles of Nir1-infected and uninfected *N. europaea* Nm50^T^ cultures. Samples were taken during the early (shortly after infection) and late (120 min) infection process as well as corresponding time points for uninfected cultures. Uninfected cultures are shown in light or dark grey, Nir1-infected cultures are shown in light or dark red, representing early or late infection time points, respectively. Values represent normalized peak areas of metabolites detected using GC-MS (n=6, except for early infected cultures, n=5), error bars represent standard deviation, n.d. = not detected. *p*-values were calculated using the Kruskal-Wallis test with Dunn’s multiple comparisons test.

## Discussion

Phages have been shown to modulate the transcriptome and metabolome of heterotrophic and phototrophic hosts to varying extent [66–69]. In this study, we extend this knowledge to viral infections of a chemolithoautotrophic microorganism involved in nitrification, a fundamental and nearly ubiquitous biogeochemical process in the N cycle [1]. We isolated Nir1 as a highly infective phage, which performs complete lysis of ammonia-oxidizing *N. europaea* Nm50^T^. Nm50^T^ is the type strain and best-studied model organism of the genus *Nitrosomonas* [25], with *Nitrosomonas* representatives being known for their presence and relevance in WWTPs [20, 23, 70]. Nir1 was detectable year-round in the WWTP Steinhof, with higher prevalence during summer months (Supplementary Fig. 1). This clearly showed that Nir1 is a stable part of the virome of this WWTP and likely followed higher population densities of its host during the warm period of the year.

The recent isolation of the related *Nitrosomonas*-infecting phage ΦNF-1 from a WWTP located in Spain [41] shows that AOB-infecting phages are likely an integral part of WWTPs. Both phages represent separate and so far the only described species of the genus *Catalonvirus*, which further underline the potential impact of this group of *Nitrosomonas*-infecting viruses in wastewaters. However, despite having >90% identical genomes, both phages exhibit different host ranges, with Nir1 infecting only *N. europaea* Nm50^T^ and ΦNF-1 infecting a broader AOB host range [41]. Electron microscopy revealed that Nir1-infected cells bloated and frequently produced membrane vesicles while characteristic intracellular membranes disintegrated (Fig. 1). This was paralleled by significantly increased concentrations of the C16-fatty acids hexadecanoate (palmitate) and hexadec-9-enoate (palmitoleate) (Fig. 5). In AOB, intracytoplasmic membranes result from invagination of the cytoplasmic membrane [27] and constitute a major structural and functional compartment, as they harbor the ammonia monooxygenase complex [71] and are highly enriched in C16 fatty acids (>96%), particularly palmitate and palmitoleate [72]. Therefore, increased levels in free C16 fatty acids are most parsimoniously explained by large-scale phospholipid hydrolysis resulting from viral takeover. Similar increases in dominant membrane fatty acids have been reported during T4 phage infection in *Escherichia coli*, reflecting breakdown of host membranes and release of lipid constituents [73, 74]. The concurrent accumulation of glycerophosphoglycerol (Fig. 5) as a major molecular backbone of phospholipids [75, 76], further supports intracellular membrane degradation by Nir1. We cannot exclude that observed increases of fatty acids and glycerophosphoglycerol relate at least in parts also to the three-step lytic release of virions in Gram-negative bacteria [77, 78]. Spanins catalyze the breakdown of the outer membrane and as such the last step in virion release during this three-step lytic process. If spanins are absent, prior endolysin-mediated degradation of the cell wall (step 2 of lytic release) can result in spherical cells [79, 80] as seen for Nm50^T^. However, in this case we should have observed pronounced trapping of viral progeny in bloated Nm50^T^ cells after infection, which was not the case (Fig. 1d).

Accumulation of fatty acids in their free form can disrupt membrane integrity, impair the respiratory electron transport chain, and inhibit membrane-bound enzymes [81]. In addition to the large-scale loss of intracytoplasmic membranes, such effects would further compromise energy conservation in Nm50^T^, given the tight coupling between ammonia oxidation, proton motive force generation, and membrane integrity. Besides ATP, also reducing power required for CO_2_ fixation by the Calvin cycle is generated by membrane-bound electron transport linked to ammonia oxidation, rendering carbon fixation highly sensitive to loss of membrane integrity as well [71, 82]. Consistent with this, Nir1 infection was accompanied by significant decreases in 3-phosphoglycerate and glyoxylate (Fig. 5). In particular, 3-phosphoglycerate represents the primary stable product of RuBisCO-mediated CO₂ fixation [83], and its depletion likely reflects collapse of Calvin cycle throughput under ATP- and NAD(P)H-limiting conditions. Glyoxylate is directly connected to the oxygenase side activity of RuBisCO. It is a typical intermediate in the phosphoglycolate salvage pathway [84], and its depletion further supports collapse of RuBisCO activity. The concurrent decline in hexose intermediates, such as fructose and galactose, aligns with these observations as it suggests failure of upstream anabolic carbon flow.

Parallel accumulation of glucose might be due to metabolic disruption of the interconnected Calvin, gluconeogenesis, oxidative and reductive pentose phosphate pathways, which share multiple enzymes and form a superpathway in *N. europaea* [85]. Such imbalance of the metabolic network may also explain increased pyruvate and succinate (central C metabolism) as well as glycolate (phosphoglycolate salvage pathway) levels during early stages of infections (Fig. 5). The reason for 3-hydroxybutanoate (3-hydroxybutyrate) accumulation is not clear. While 3-hydroxybutanoate is the well-established monomer of the storage compound polyhydroxybutyrate (PHB) [86], PHB accumulation in *Nitrosomonas* species has not been reported as to the best of our knowledge. PHB may have transiently formed in our experiments and used as carbon source by the host during Calvin cycle breakdown. Alternatively, 3-hydroxybutanoate accumulated because of metabolic disruption by Nir1, caused by incomplete processing of acetyl-CoA and reducing equivalents to be channeled towards 3-hydroxybutanoate as a sink for excess carbon and reducing equivalents [86].

Viral infection also caused altered profiles of free amino acids, although changes were not very pronounced (Fig. 5). This is likely explained by the similar amino acid composition of the overall host proteome as compared to the overall phage proteome or even the amino acid composition of its major capsid protein (Supplementary Table 2). Intriguingly, the arginine-derived diamine putrescine was highly upregulated during the late infection process (Fig. 5). Putrescine represents next to spermidine and spermine one of the major polyamines in bacteria. All three are small cationic compounds known to bind DNA and were shown before to be beneficial for phage replication and genome packaging [69, 87, 88]. However, the role of putrescine can be ambivalent as it was also shown to support anti-phage responses in *Pseudomonas aeruginosa*. Here, putrescine served as danger signal released by lysed cells. Adjacent host cells were actively taking up released putrescine, where it interfered with phage replication [89].

Parallel host transcriptome profiling identified overexpression of genes involved in transcriptional regulation (RNA polymerase sigma factors) and translation (ribosomal proteins) as well as general stress responses and DNA synthesis, replication and repair (Fig. 4, Supplementary Table 1). Such responses are also known from the infection process of other lytic phages [90, 91]. In addition, genes involved in starvation response or nutrient uptake were overexpressed in Nir1-infected Nm50^T^ during the late infection process. This effect was very pronounced for genes associated with iron scavenging. *N. europaea* is known to have a high iron demand, mainly for heme and Fe-S cluster containing enzymes of its respiratory chain [92, 93]. Also, under iron-replete conditions *N. europaea* forms less intracytoplasmic membrane layers [93]. Since Nir1-infection caused a massive breakdown of intracytoplasmic membranes, scavenging of iron may have supported synthesis of iron-containing respiratory chain proteins to counterbalance production loss of ATP and reducing equivalents. This would benefit the replicating virus and prolong the lifespan of the disintegrating host cell. In line with this observation is the overexpression of genes encoding proteins involved in electron transfer, such as cytochrome c and multicopper oxidases (Supplementary Table 1). The latter have been proposed to be involved in the respiratory chain in ammonia oxidizing archaea, possibly substituting for iron-containing cytochromes [94]. A starvation response is further indicated by overexpression of one *cstA-* and one *ybdD-*related upregulated gene in late infected cells (Supplementary Table 1). Both have been shown to play roles in the uptake of pyruvate in carbon-starved cells of *Escherichia coli* [95]. Previous studies revealed that pyruvate [96] and amino acids [97, 98] can support growth of *N. europaea* cultures and that chemolithoorganotrophic growth on organic compounds, including pyruvate [28], is possible.

We observed that infections with Nir1 resulted in the overexpression of flagellar and type IV pilus genes in the late infection process. Interestingly, genes encoding the actual filament subunits (*flaA* and *pilA*) were only slightly overexpressed (LFC 0.33 and 0.57, respectively) in comparison to other genes of the flagella and pilus apparatus. This is supported by our electron microscopy investigation which did not reveal a massive switch to a flagellated phenotype of Nm50^T^. Type IV pili and flagella can play multiple roles for bacteria and their associated phages and are best known for their potential to facilitate motility, via rotation or by twitching motility, respectively. However, pores of type IV pili can also play a role in the uptake of extracellular DNA, which involves presence of the highly conserved ComEC protein to enable transport through the cytoplasmic membrane [99]. We found a highly overexpressed gene coding for a ComEC/Rec2 type protein in the late infection process, indicating that DNA-uptake might play a role in Nir1-infected cells of Nm50^T^.

In addition, we detected the upregulation of multiple genes involved in the synthesis of nucleotides, further supporting an elevated demand of DNA to produce progeny virions. The parallel uptake of external amino acids was indicated by the overexpression of genes encoding an amino acid permease (Supplementary Table 1). Furthermore, we observed a strong overexpression of genes coding for a urea carboxylase and urea amidolyase-related genes (Fig. 4, Supplementary Table 1). Interestingly, Nm50^T^ lacks genes encoding the classical urease [92]. Urea carboxylase is part of the urea amidolyase complex and can be used as alternative to urease for the conversion of urea into ammonia and CO_2_ [100]. However, *N. europaea* has not been reported to grow on urea and urea carboxylase can also serve as component of a pyrimidine nucleic acid precursor degradation pathway [100]. We regard this as a more likely function in Nir1-infected Nm50^T^ considering phage-induced intracellular N salvage and redistribution to meet the high N demand of viral progeny [101].

## Conclusions

Our study provides the first detailed description of phage-induced host modulation of a chemolithoautotrophic ammonia oxidizing bacterium. We show that virus-induced intracytoplasmic membrane disintegration simultaneously compromised lipid homeostasis, energy conservation, and autotrophic carbon metabolism prior to cell lysis. To counterbalance the demand for reducing equivalents and ATP an upregulation of the iron-dependent respiratory chain was indicated to support the enzymatic machinery in the remaining membrane of bloated cells. In parallel, uptake systems for nucleic acids, amino acids but also small organic compounds such as pyruvate likely supported metabolic rewiring of the host to support build-up of viral progeny. Our results are important in three ways. Since Nir1 infects *Nitrosomonas europaea* as one of the best studied AOB, the Nir1-Nm50^T^ phage-host system may serve in future as an important model to understand the biology of AOB-infecting phages. Furthermore, our results provide an important basis and first step to assess the impact phages can have on the performance of WWTPs, especially in the background of highly efficient lysis, varying host ranges, and lipid release. Since AOB play also an important role in agricultural systems, leading to fertilizer loss [2] and emissions of the greenhouse gas N_2_O [3–5], the presented results will also be important to assess the potential application of phage cocktails for environmental nitrification control. Future research needs to resolve the molecular mechanisms governing phage–AOB interactions and to quantify the environmental consequences of AOB-infecting viruses across engineered and natural ecosystems.

## Acknowledgements

We are thankful to the operators of the WWTP Steinhof for providing samples. We thank Stefan Dyksma for support in bioinformatics.

## Funding

Funding was provided to JP and MP by the German Research Foundation within grant PE2147/5-1.

## Data availability

The sequenced genome of *Nitrosomonas europaea* Nm50^T^ (=DSM 28437) has been deposited at DDBJ/ENA/GenBank under the accession PRJNA1375795. The sequenced genome of vB_NeuP-Nir1 has been deposited at DDBJ/ENA/GenBank under the accession PX682272. Transcriptomic sequencing data has been deposited at NCBI SRA under the accession PRJNA1380931. Metabolomic data has been deposited at Fairdomhub.

## Notes

### Competing Interest Statement

The authors have declared no competing interest.

