## Supplementary Material for "Viral infection collapses intracytoplasmic membrane integrity and autotrophic metabolism in ammonia-oxidizing *Nitrosomonas europaea*"

### Supplementary Material and Methods

Composition DSMZ medium 1583

The main solution of DSMZ medium 1583 is composed of 535 mg $L^{-1}$ $NH_4Cl$, 54 mg $L^{-1}$ $KH_2PO$, 74 mg $L^{-1}$ KCl, 49 mg $L^{-1}$ $MgSO_4 \times 7H_2O$, 147 mg $L^{-1}$ $CaCl_2 \times 2H_2O$ and 584 mg $L^{-1}$ NaCl in distilled deionized water. The medium was autoclaved and subsequently amended with 2 mL $L^{-1}$ autoclaved and 0.2-µm-filtered (Filtropur S 0.2, PES, Sarstedt, Nümbrecht, Germany) cresol red solution (0.5 g $L^{-1}$ in distilled deionized water) and 1 mL $L^{-1}$ autoclaved trace element solution. The latter comprises 975 mL distilled deionized water amended with 25 mL 1M HCl, 45 mg $L^{-1}$ $MnSO_4 \times 4H_2O$, 49 mg $L^{-1}$ $H_3BO_3$, 43 mg $L^{-1}$ $ZnSO_4 \times 7H_2O$, 37 mg $L^{-1}$ $(NH_4)_6Mo_7O_{24} \times 4H_2O$, 973 mg $L^{-1}$ $FeSO_4 \times 7H_2O$ and 25 mg $L^{-1}$ $CuSO_4 \times 5H_2O$. The pH of DSMZ medium 1583 was initially adjusted by slow addition of autoclaved 10% w/v $NaHCO_3$ until the color of the pH indicator changed from yellow to pink at pH ~7.8. pH had to be regularly re-adjusted by addition of autoclaved 10% w/V $NaHCO_3$ solution during bacterial growth.

Staining and counting of virions

Staining of virions was performed according to a technique described by Kim, Kim (1) with minor modifications, as described in the following. Stainless steel Luer lock syringe filter holders were equipped with a 0.45 µm cellulose nitrate support filter (Sartorius, Göttingen, Germany) and a 0.02 µm aluminum oxide (Whatman™ Anodisc™, Global Life Sciences Solutions Operations UK Ltd., Little Chalfont, UK) filter to capture virions. Viral lysate in the host medium (DSMZ medium 1583) was pre-filtered through 0.2 µm PES filters (Filtropur S 0.2, Sarstedt) to remove host debris. Thereafter, 1 mL of pre-filtered viral lysate was further filtered through the 0.02 µm aluminum oxide filter in the assembled stainless-steel filter capsule (see above) using a 2.5 mL syringe. As a final step, 1 mL of sterile 0.2-µm-filtered DSMZ medium 1583 was used to flush the syringe and the 0.02 µm aluminum oxide filter. The aluminum oxide filters were then immersed in 0.1% low melting Agarose (TopVision, Thermo Fisher Scientific Baltics UAB) to minimize dislodging of virions from the membrane during staining and observation. Thereafter, filters were transferred to a microscope slide. Staining of virions was performed by pipetting 100 µL of 1:10,000 diluted SYBR Gold (in TE Buffer) on the filter followed by incubation for 10 min in the dark. Afterwards, the stain was carefully pipetted off. To reduce bleaching of stained virions during observation, the filter was covered using 20 µL of a 4:1 mixture of Citifluor AF1 (Electron Microscopy Sciences, Hatfield, USA) and VECTASHIELD (Vector Laboratories, Burlingame, USA). Investigation was done immediately after staining using a BZ-X810 Microscope (KEYENCE DEUTSCHLAND GmbH, Neu-Isenburg, Germany) equipped with a 470/40 nm excitation and 525/50 nm emission filter set. Pictures of the stained virions were captured using the supplied software package BZ-X Analyzer. Calculation of virions concentration was performed by adding a counting grid of 10×10 µm or 20×20 µm and extrapolating the results of the filtered volume.

Isolation of genomic DNA and reconstruction of the closed genome of *N. europaea* Nm50$^T$

DNA of strain Nm50$^T$ was isolated from bacterial biomass collected by centrifugation (9,000×*g*, 30 min, 4°C) using the MasterPure™ Complete DNA & RNA Purification Kit (Lucigen Corp., Middleton, USA). DNA extraction was performed following the manufacturer's instructions for cell samples but using double the volumes mentioned in the protocol. In brief, 2 µL Proteinase K solution were diluted in 600 µL of Tissue and Cell Lysis Solution and added to the bacterial pellet. The sample was mixed thoroughly on a vortex mixer and incubated at 65 C° for 15 min. During incubation, the sample was vortexed every 5 min. This was followed by cooling to 37C° for 5 min, adding 2 µL of 5 mg mL$^{-1}$ RNase A, thorough mixing and incubation at 37C° for 30 min. Thereafter, the sample was placed on ice for 5 min. Afterwards, 300 µL of MPC Protein Precipitation Reagent were added to 600 µL of lysed sample and mixed thoroughly using a vortex mixer. The debris was pelleted by centrifugation at 4 C° for 10 minutes at 10,000×*g*. The supernatant was transferred to a clean 2 mL microcentrifuge tube, discarding the pellet. The supernatant was amended with 1000 µl of isopropanol followed by mixing by repeatedly inverting the microcentrifuge tube. The DNA was then pelleted by centrifugation at 4 C° for 20 min at 20,817×*g*. The isopropanol was carefully poured off without dislodging the DNA pellet which was then rinsed with freshly prepared 70 % ethanol. The residual ethanol was removed with a pipette and the pellet was air-dried. Finally, the DNA was resuspended in 50 µL TE buffer.

The genome of Nm50$^T$ was long-read sequenced on a PacBio Revio system (Pacific Biosciences, Menlo Park, USA) and assembled using the Microbial Genome Analysis Protocol within SMRTLink v13.1. Annotation of the genome was performed using the NCBI prokaryotic genome annotation pipeline [2].

### Supplementary Results

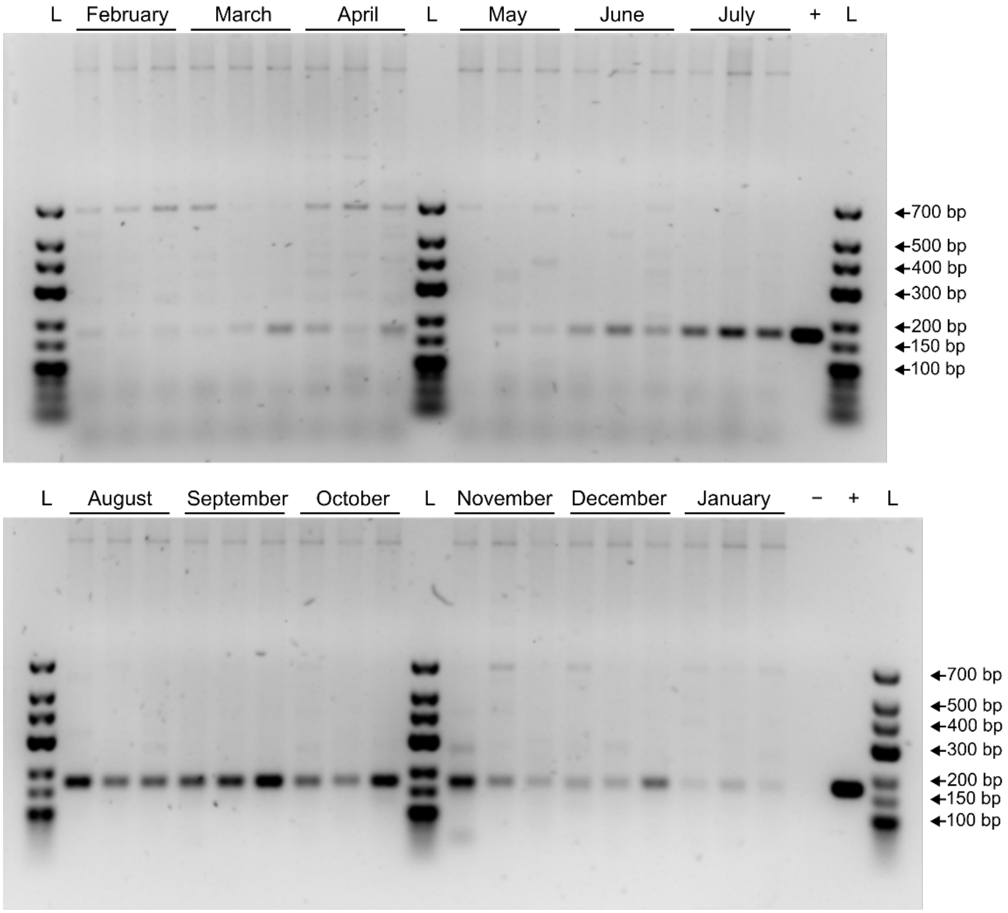


**Supplementary Figure 1.** Products of PCR-amplified 170 bp long region of the major head protein of phage vB_NeuP-Nir1, separated by agarose gel electrophoresis using 2% Wide Range agarose (SERVA Electrophoresis GmbH, Heidelberg, Germany) in 1× TAE buffer (Carl Roth GmbH + Co. KG, Karlsruhe, Germany). Negative control (-) = nuclease-free water (Thermo Fisher Scientific Baltics UAB, Vilnius, Lithuania), positive control (+) = 0.121 ng genomic DNA of phage vB_NeuP-Nir1, Ladder (L) = GeneRuler Low Range DNA Ladder (Thermo Fisher Scientific Baltics UAB).

**Supplementary Table 1.** Differentially expressed genes of Nir1-infected *N. europaea* Nm50$^T$ compared to non-infected controls during the late infection process (sampled 120 min post infection). Genes are marked according to their assigned functional category. (**Separate Excel file**)

**Supplementary Table 2.** Comparison of amino acid (AA) frequencies in total CDS of the host bacterium *N. europaea* Nm50$^T$, total CDS of the infecting bacteriophage vB_NeuP-Nir1 (Nir1) and its major head protein.

| Amino acid | Amino acid single letter | AA frequency in Nm50$^T$ CDS | AA frequency in Nir1 CDS | AA frequency in Nir1 major head protein |
|---|---|---|---|---|
| Alanine | A | 9.20% | 7.90% | 10.80% |
| Cysteine | C | 1.00% | 0.90% | 0.30% |
| Aspartate | D | 5.30% | 5.90% | 7.00% |
| Glutamate | E | 6.00% | 6.10% | 4.40% |
| Phenylalanine | F | 3.90% | 3.20% | 4.40% |
| Glycine | G | 7.20% | 7.60% | 7.30% |
| Histidine | H | 2.50% | 1.90% | 2.60% |
| Isoleucine | I | 6.30% | 5.50% | 4.40% |
| Lysine | K | 4.10% | 5.60% | 4.70% |
| Leucine | L | 10.60% | 8.50% | 7.90% |
| Methionine | M | 2.40% | 3.10% | 2.60% |
| Asparagine | N | 3.60% | 4.50% | 4.70% |
| Proline | P | 4.60% | 4.00% | 3.20% |
| Glutamine | Q | 4.30% | 3.70% | 3.80% |
| Arginine | R | 6.40% | 5.60% | 5.60% |
| Serine | S | 6.00% | 7.40% | 6.10% |
| Threonine | T | 5.50% | 5.70% | 8.50% |
| Valine | V | 6.70% | 7.10% | 6.70% |
| Tryptophan | W | 1.30% | 1.50% | 2.00% |
| Tyrosine | Y | 2.90% | 3.90% | 2.60% |
| STOP | * | 0.30% | 0.50% | 0.30% |
